# Cellular memory enhances bacterial chemotactic navigation in rugged environments

**DOI:** 10.1101/733345

**Authors:** Adam Gosztolai, Mauricio Barahona

**Affiliations:** Department of Mathematics, Imperial College London, SW7 2AZ, London, United Kingdom

## Abstract

The response of microbes to external signals is mediated by biochemical networks with intrinsic time scales. These time scales give rise to a memory that impacts cellular behaviour. Here we study theoretically the role of cellular memory in *Escherichia coli* chemotaxis. Using an agent-based model, we show that cells with memory navigating rugged chemoattractant landscapes can enhance their drift speed by extracting information from environmental correlations. Maximal advantage is achieved when the memory is comparable to the time scale of fluctuations as perceived during swimming. We derive an analytical approximation for the drift velocity in rugged landscapes that explains the enhanced velocity, and recovers standard Keller-Segel gradient-sensing results in the limits when memory and fluctuation time scales are well separated. Our numerics also show that cellular memory can induce bet-hedging at the population level resulting in long-lived multi-modal distributions in heterogeneous landscapes.

## INTRODUCTION

The natural habitat of many microbes is shaped by inherent micro-scale ruggedness arising from random spatial inhomogeneities due to porous or particulate matter^1–3^) or to filamentous structures resulting from turbulent advection in the medium^4,5^. Microbes typically navigate such rugged attractant landscapes in search of nutrients and stimulants in a process called chemotaxis. Chemotaxis is mediated and governed by specialised biochemical pathways that sense changes in stimulant concentration, transduce those signals, and induce subsequent adjustments to the locomotion of the cell^6^. Such pathways have characteristic dynamic responses with intrinsic time scales, which are used by cells to resolve changes in chemoattractant concentrations, i.e., to perform local gradient-sensing^7,8^. In addition, the dynamic response of the biochemical circuits can filter out the high frequencies of noisy signals, to enhance gradient-sensing^9–11^.

The time scales of such responses can also be viewed as the basis for a cellular memory, over which signals are processed. Indeed, microbes sample continuously their chemical environment along their swimming trajectory, and recent work has shown that the biochemical memory can be dynamically tuned^12^ from seconds to minutes^13^ in response to environmental statistics. Hence, in addition to evaluating the stimulant gradient, cells could extract informative features of the heterogeneous environment from the fluctuations they perceive as they swim.

We study the effects of cellular memory in the context of *Escherichia coli* (*E. coli*) chemotaxis, a model system for the navigation of microbes^5^, worms^14^, and eukaryotes^15^, as well as an inspiration for the motion of swarm robots^16,17^ and random search algorithms^18^. *E. coli* chemotaxis entails a run-and-tumble strategy: runs (i.e., stretches of linear motion at constant velocity) interrupted by tumbles (i.e., random stops with reorientation onto a random direction). To generate a drift towards high chemoattractant concentrations, cells reduce their tumbling rate upon sensing a favourable gradient, thus lengthening the up-gradient runs^19^.

The tumble rate is regulated by a chemotactic pathway with a bi-lobed temporal response with a characteristic time scale *γ*, which we denote the cellular memory. Input signals are convolved with this temporal response, with the effect that recent samples are weighed positively whereas signals in the past are given a negative weighting^20^. It has been shown that this response yields an estimate of the local temporal gradient^8,9^.

The capability of cells to compute local gradients is the basis for several coarse-grained models (drift-diffusion equations). The classic example is the linear Keller-Segel (KS) model^21,22^, which describes the behaviour of a population of cells whose mean velocity aligns instantaneously with the local gradient. The KS model successfully reproduces a variety of chemotactic phenomena, including experimentally observed distributions under shallow attractant gradients^23^. Yet the presence of fluctuations may lead to an incorrect assessment of the underlying gradient if using only instantaneous information. Indeed, KS fails to recapitulate situations when cells do not have time to adapt to large fluctuations, both in experiments^24^ and in agent-based simulations^25–27^. These shortcomings suggest the need to consider additional time scales that play a role in chemotactic transient responses^28^, and, specifically, the intrinsic memory of the chemotactic pathway processing incoming stimuli^29,30^.

Here, we study how bacteria use their cellular memory as they swim across a rugged chemoattractant landscape to extract spatio-temporal information from the perceived signal to improve their chemotactic navigation. To shed light on the role of memory, we carry out simulations of an agent-based (AB) model containing an input-output response function of the *E. coli* chemotactic pathway^31–33^ and compare its predictions to the KS model, which is based on memoryless local gradient alignment. The KS agrees well with the AB numerics for constant gradients, yet it underestimates the drift velocity of the population when the ambient concentration has spatial correlations, consistent with cells taking advantage of correlations in addition to local gradients.

Motivated by these numerical findings, we derive an analytical formula for the drift velocity in terms of the cellular memory and the length scale of the spatial correlations of the attractant landscape. Our model predicts the numerical results and recovers KS in various limits, thus elucidating the conditions in which cellular memory provides a chemotactic advantage over memoryless local gradient-sensing. We also show that our results are consistent with optimal information coding by the chemotaxis pathway^32,34^, yet cells are band-limited by their tumbling rate. Our work thus extends the gradient-sensing viewpoint in chemotaxis, and provides insight into the role of memory in navigating heterogeneous landscapes.

## RESULTS

### GRADIENT-SENSING AS THE CLASSICAL VIEWPOINT OF CHEMOTAXIS

A classical setup for chemotaxis is represented schematically in Fig. 1a. Cells swim following a run-and-tumble motion: ballistic motion (‘runs’) at constant velocity *v*_0_, interrupted by random re-orientations (‘tumbles’) occurring at random times governed by a Poisson process with rate λ(*t*)^35^. As cells swim along their trajectory *x*(*t*) (taken here to be one-dimensional for simplicity), they are exposed to an attractant concentration *S*(*x*(*t*)). Assuming initial adaptation to the ambient attractant concentration, the cells modulate their Poisson tumbling rate according to^29,36^:

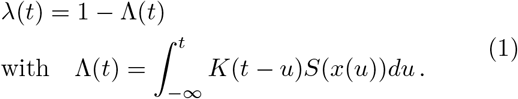

**Figure 1.**
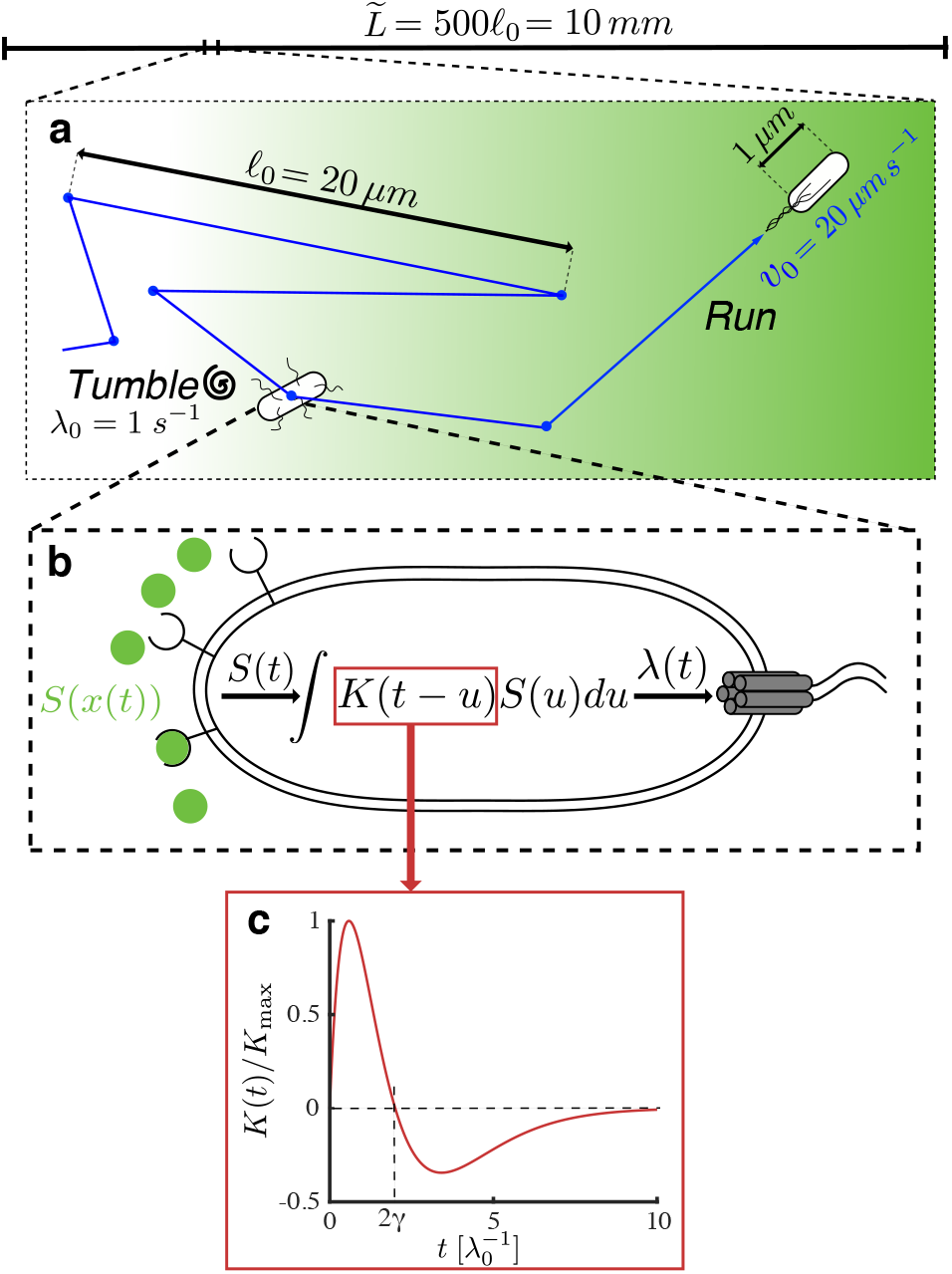
Setup of the agent-based model and simulation framework. **a** Cells navigate a chemoattractant landscape *S*(*x*) using a run and tumble strategy with characteristic scales and variables as represented in the picture (*ℓ*_0_, 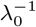 and *v*_0_ are the typical run length, run time and ballistic run speed respectively). The simulations are run in a long domain of length 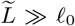 over long times 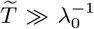. **b** The swimming cell senses the attractant concentration along its trajectory *x*(*t*) and modulates its tumbling rate λ(*S*(*x*(*t*))) by the chemotaxis transduction pathway with response given by Eq. (1). The dynamic response of the pathway is mediated by the response kernel *K*(*t*) (Eq. (2)). **c** The shape of the bi-lobed kernel *K*(*t*) normalised by its amplitude *K*_max_ against time.

This represents the dynamics of the chemotaxis signal transduction, illustrated on Fig. 1b. Throughout, we use variables non-dimensionalised with respect to the characteristic length and time scales:

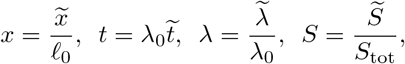

where *S*_tot_ is the total attractant concentration, λ_0_ = 1 *s*^−1^ is the basal tumbling rate, and *ℓ*_0_ = *v*_0_/λ_0_ = 10 *μm* is the average run length^19^.

In Eq. (1), *K*(*t*) is the chemotactic memory kernel, measured through impulse response experiments^20^, which has a bi-lobed shape for some attractants in *E. coli*^37,38^ (Fig. 1c). A typical form for *K*(*t*) is given by:

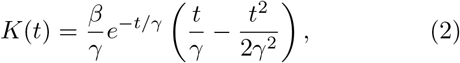

where *β* is a dimensionless signal gain, and 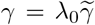 is the cellular memory, a (dimensionless) relaxation time, as seen by the fact that the crossing point of the bi-lobed response is *t* = 2*γ* (Fig. 1c). Note that the amplitude of the response kernel is 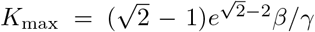. Hence an increase in memory decreases the overall response. The kernel in Eq. (2) can be understood as a linear filter with three states (Supplementary Note 1), with a topology that achieves perfect adaptation^39,40^, since 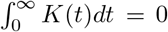. Our setup differs from an alternative model^26^ based on the dynamics of the CheY protein regulating the tumbling rate, which typically leads to a one-state linear filter obtained through linearisation. In Supplementary Note 2, we show that this alternative model can be equivalently written in the form of Eq. (1) but with a kernel *K*(*t*) = *β*(*δ*(*t*) − (1/*γ*)*e*^−*t*/*γ*^) with a singularity at *t* =0 and a single decaying exponential, instead of the bi-lobed kernel (Eq. (2)).

At long time scales involving many runs and tumbles (*t* ≫ 1), the swimming behaviour may be approximated by a drift-diffusion process^22,25,41^. In this regime, the time evolution of the population density of cells *ρ*(*x, t*) from an initial state *ρ*(*x*, 0) is described by a Fokker-Planck partial differential equation:

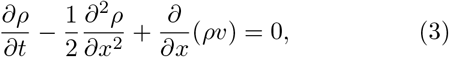

where 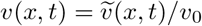 is the drift velocity of the cells, and the diffusion coefficient 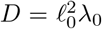 drops out as part of the non-dimensionalisation. Equivalently, *ρ*(*x*, *t*) is the probability of finding a cell at *x* after time *t* from a starting position *x*_0_ drawn from *ρ*(*x*, 0).

Typically, derivations in the literature^35,41,42^ consider the regime of long memory (compared to the average run) and shallow perceived gradient (i.e., the attractant does not vary appreciably over the memory):

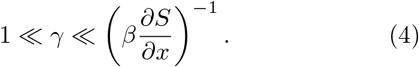

Under these assumptions, the drift velocity *v*(*x, t*) can be shown^36^to align with the local gradient (see Supplementary Note 3):

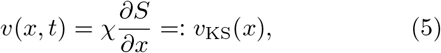

where the chemotactic response coefficient *χ* follows from the kinematics and the memory kernel (Eq. (2)):

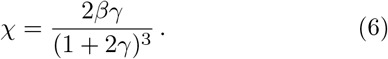

Eq. (3), together with Eqs. (5) and (6), defines the classic linear Keller-Segel (KS) equation for the time evolution of the population density under a landscape *S*(*x*). We denote the solution to this equation as *ρ_KS_*(*x*, *t*; *S*).

However, the KS model is actually valid under the weaker condition^25^

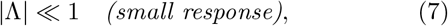

i.e., the tumbling response remains close to the adapted value. It can be shown that time scale separation (Eq. (4)) implies small response (Eq. (7)), but the converse is not necessarily true. Hence KS can still be valid in the realistic situation when Eq. (4) breaks down because the cellular memory is commensurate with environmental fluctuations^10,11,32^, as long as Eq. (7) holds. Below, we consider a broad span of memory values (from the well separated to the commensurate) but always in the small response regime so that the KS model is valid.

### CHEMOTAXIS OF CELLS WITH MEMORY STUDIED USING AGENT-BASED NUMERICS

We consider cells with memory swimming in a rugged environment with spatial correlations, leading to a temporally fluctuating input perceived along their trajectories. To study the effect of memory, we performed agent-based (AB) simulations of run-and-tumble motion as in Refs.^31,32^ coupled to a cellular response (Eqs. (1)–(2)) with memory (see Supplementary Note 1 for details).

Our rugged landscape is a simple linear attractant concentration profile with additive spatial noise^32^:

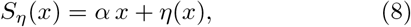

where *η*(*x*) is a random spatial variable described by the stochastic harmonic oscillator Langevin equation:

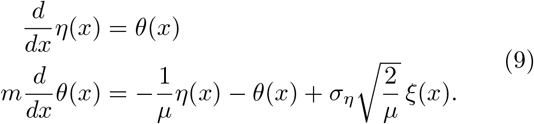

Here *ξ*(*x*) is a unit white noise, and

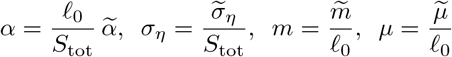

are non-dimensionalised parameters corresponding to: attractant gradient, noise variance, inertia, and spatial correlation length, respectively. This random landscape has two desirable properties. First, *S_η_*(*x*) with *m* > 0 is continuous and differentiable, so that

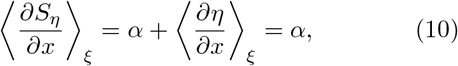

where 〈·〉_*ξ*_ denotes averaging over independent realisations of *η*(*x*). Second, Eq. (9) is a regularised spatial Ornstein-Uhlenbeck (OU) process (Supplementary Figure 1): as *m* → 0, *η* converges to an OU process *η*^0^(*x*) which has exponential correlations with characteristic length *μ*:

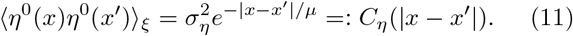

The OU limit is used below to facilitate our analytical calculations.

### CHEMOTAXIS IN CONSTANT SHALLOW GRADIENTS

We first consider the landscape with zero ruggedness:

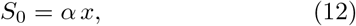

which corresponds to *σ_η_* = 0 or, alternatively, to the limit *μ* → ∞, when the correlation length diverges. In this case, it has been shown^22,42^ that the condition

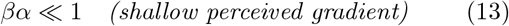

guarantees that the small response condition (Eq. (7)) also holds. Hence we expect the AB numerics to be well described by the KS equation.

To test this prediction, we used the AB model to simulate *N* = 10^5^ independently generated cell trajectories {*x*_AB_(*t*; *S*_0_); *t* ∈ (0, *T*)}, where *T* = 4 × 10^3^ (Fig. 2a), from which we obtain population snapshots, *ρ_AB_*(*x, t*; *S*_0_). All the simulations were run in the regime of small *βα*. In Fig. 2b, we show that the statistics of the AB simulations are well captured by the continuum KS solution:

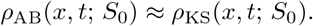

**Figure 2.**
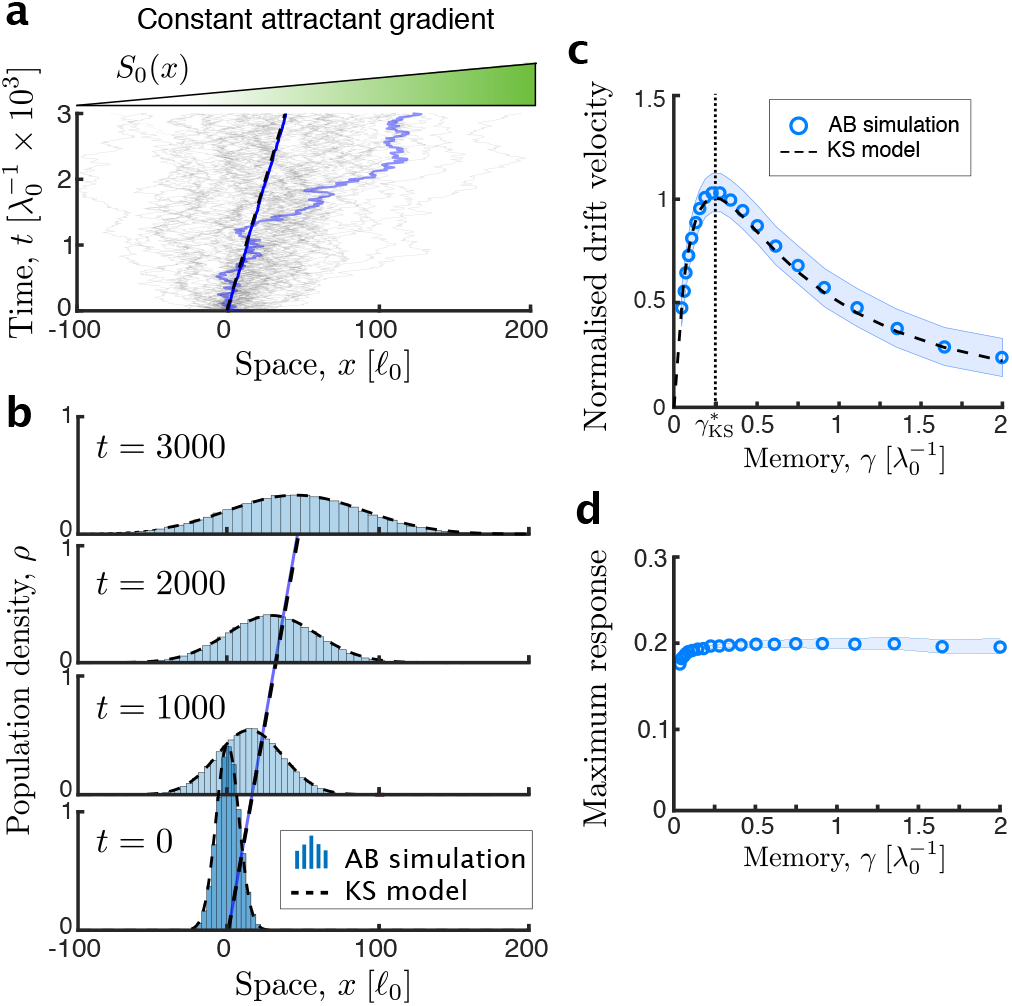
Comparison of agent-based numerics and Keller-Segel approximation in shallow gradients. The AB model is used to produce *N* = 10^5^ cell trajectories over *T* = 4 × 10^3^ (Δ*x* = 5 × 10^−5^, Δ*t* = 5 × 10^−3^) with perceived gradient *βα* = 0.1, *βσ_η_* = 0. The KS model is integrated numerically using a first-order in time, second-order in space forward-Euler scheme (Δ*x* = 10^−4^, Δ*t* = 1). **a** Sample trajectories of the AB model (*σ* = 0.5) in the deterministic landscape *S*_0_(*x*). **b** The evolution of the population density of the AB model (*ρ*_AB_(*x*, *t*; *S*_0_), histogram) is well captured by the evolution of the KS equation (*ρ*_KS_(*x*, *t*; *S*_0_), Eq. (3), dashed line, *σ* = 0.5). The solid blue line indicates the average velocity vab, which is indistinguishable from the KS drift *v*_KS_. The densities are normalised to unit mass. **c** The drift velocity from the AB model (*v*_AB_, circles) is well predicted by the KS drift velocity (*v*_KS_, dashed line). Both velocities are normalised by the maximal KS drift velocity 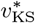, which is reached at a memory of 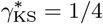, shown by the dotted line. **d** The maximum tumbling response max |Λ| (circles) stays well below unity, showing that the small response condition is met for all the simulations. The blue band indicates the standard deviation of the simulations.

We also compared the drift velocity of the KS solution to the average velocity of AB cells (computed over the long simulation time *T*):

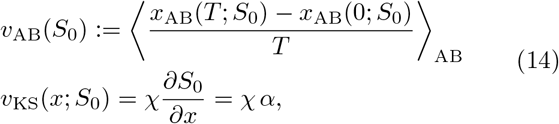

where 〈·〉_AB_ denotes averaging over the ensemble of AB cells. Fig. 2c shows that the average velocity of the AB population matches the drift velocity of the KS model for varying memory *γ*.

Maximising Eq. (6) shows that the drift velocity *v*_KS_ achieves a maximum at an optimal memory:

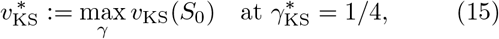

Hence a cell with optimal memory 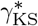 has a kernel *K*(*t*) with a zero crossing at *t* = 1/2, i.e., halfway through the expected length of a run (see Fig. 1c). KS thus predicts that the drift speed is maximal when the gradient is measured along a single run, when the cell can take an unbiased measurement while moving in a straight line. For the zero-ruggedness landscape, our AB simulations (Fig. 2c) also display a maximum in the average velocity of the population when the cells have memory 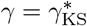.

Fig. 2d confirms that the simulations are in the regime of small response (Eq. (7)) where KS holds. As *βα* is increased, and the small response condition (Eq. (7)) is violated, the correspondence between the AB and KS solutions gradually breaks (see Supplementary Figure 2).

### CHEMOTAXIS IN RUGGED, CORRELATED LANDSCAPES

The kernel *K*(*t*) with intrinsic memory *γ* has been shown to filter high-frequency input noise^11,32^. However, cells could also use this memory to their advantage as they process the correlated fluctuations that they encounter as they traverse a rugged landscape.

To test this idea, we carried out AB simulations of cells with memory navigating the spatially correlated landscape (Eq. (8)) and compared it to the predicted KS behaviour. To ensure that the differences between AB and KS are a direct consequence of the correlated spatial fluctuations, all our simulations are run in the small response regime (Eq. (7)) where KS holds, while keeping a large signal-to-noise ratio (Supplementary Note 4):

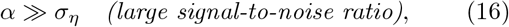

Fig. 3 shows that the AB cell population travels faster than predicted by KS going up the gradient of the rugged landscape. Fig. 3 a presents simulated AB trajectories for a particular realisation of the landscape *S_η_*(*x*), and Fig. 3b compares the time evolution of the KS solution 〈*ρ*_KS_(*x*, *t*; *S_η_*)〉_*ξ*_ to the empirical distribution from the AB numerics 〈*ρ*_AB_(*x*, *t*; *S_η_*)〉_*ξ*_, both averaged over 10^2^ independent realisations of the landscape *S_η_*(*x*). Our numerics show that the AB distribution propagates faster: 〈*v*_AB_(*S_η_*)〉_*ξ*_ > *v*_KS_, i.e., the average cell velocity of the AB simulations (defined in Eq. (14)) averaged over realisations of the landscape (blue solid line) is larger than the corresponding KS drift velocity (dashed line).

**Figure 3.**
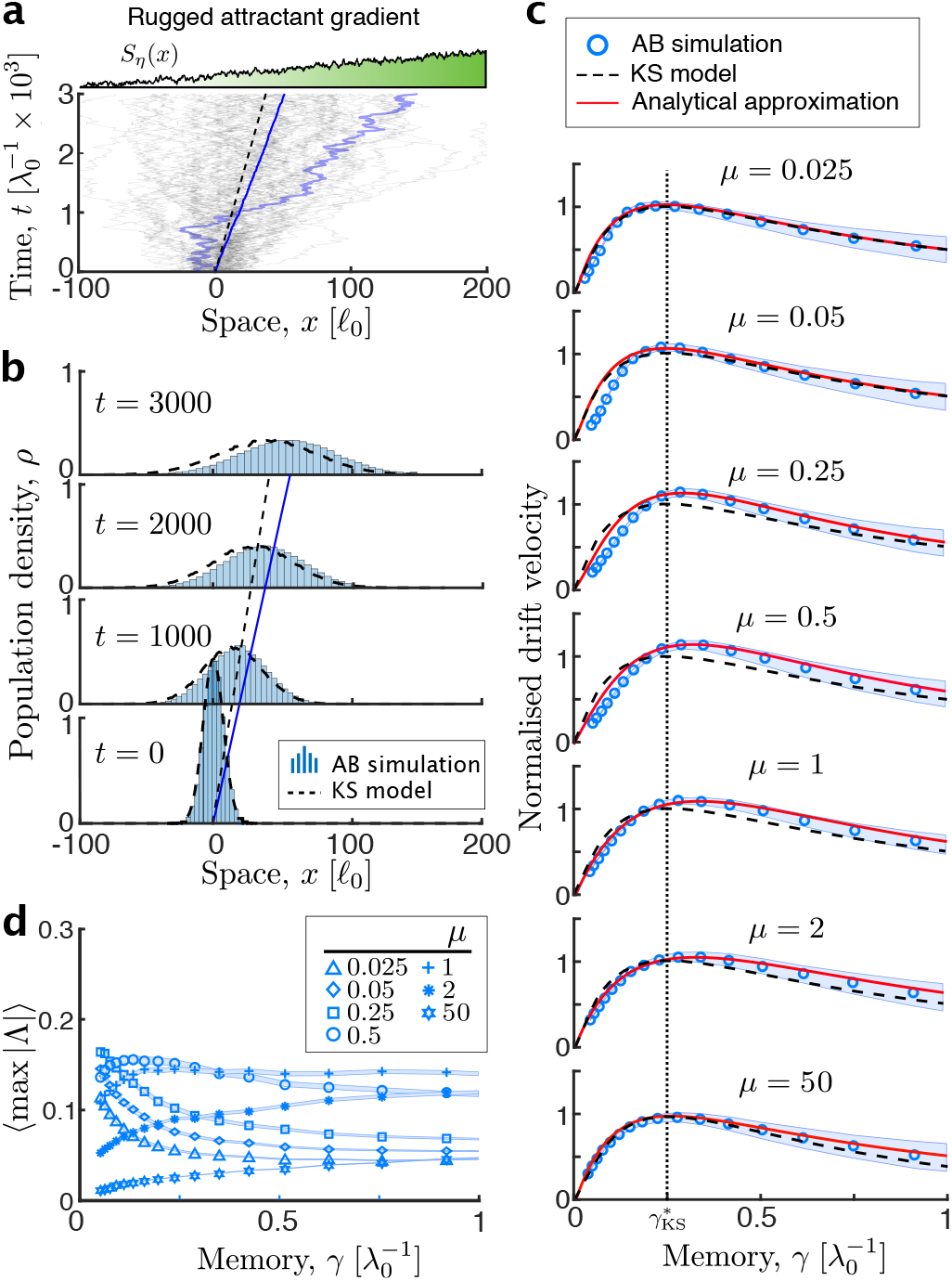
Comparison of agent-based simulations and analytic predictions in rugged landscapes *S_η_*. The AB model is used to produce *N* = 10^4^ cell trajectories over *T* = 4 × 10^3^ (Δ*x* = 5 × 10^−5^, Δ*t* = 5 × 10^−3^) in 10^2^ realisations of *S_η_* (*x*) with perceived gradient *βα* = 0.05, *βσ_η_* = 10^−3^. The KS model is integrated numerically using a first-order in time, second-order in space forward-Euler scheme (Δ*x* = 10^−4^, Δ*t* = 1). **a** Sample trajectories of the AB model (*γ* = 0.5) in the rugged landscape (*μ* = 1). Note that the ensemble average drift of the AB cells (solid blue line) is faster than the KS drift (dashed black line). **b** The evolution of the population density of the KS model (〈*ρ*_KS_(*x,t; S_η_*)〉_*ξ*_, Eq. (3), dashed black line, *γ* = 0.5, *m* = 5 × 10^−3^) generally fails to capture the evolution of the AB population (〈*ρ*_AB_(*x, t; S_η_*)〉_*ξ*_, histogram). The densities are normalised to unit mass. **c** The average drift velocity of the AB model (〈*v*_AB_(*S_η_*)〉_*ξ*_, circles) as a function of the memory *γ* is well described by our approximation (*V_μ_*, Eq. (23), red solid line) for all values of *μ*, but in general not well captured by the KS drift (*v*_KS_, Eq. (5), black dashed line). The AB drift is averaged over realisations of the rugged landscape and both velocities are normalised by the maximal KS drift 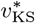 in the deterministic landscape. **d** The average maximum response amplitude |Λ| from the AB simulations at different values of *μ* (different symbols in the figure) remains much smaller than unity for all combinations of *γ* and *μ* (as in Fig. 2d), indicating that the KS model still holds in the rugged landscape. Yet, as panels b, c show, it does not capture the drift velocity. The blue bands indicate the standard deviation of the simulations.

We have examined this enhanced chemotaxis as a function of the length scale of the landscape. We show in Fig. 3c that for correlation lengths around the run length (*μ* ≈ 1), the KS drift velocity *v*_KS_ underestimates the average velocity 〈*v*_AB_(*S_η_*)〉_*ξ*_ of cells with memory 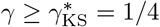.

As expected, the average velocity of the AB cells is well approximated by the KS solution in the limits of both vanishingly small and infinitely large correlation length:

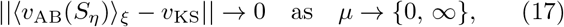

which correspond to an uncorrelated landscape or a zeroruggedness constant gradient, respectively. Also expected, Eq. (17) holds in the memoryless limit *γ* → 0. In this limit, the kernel *K*(*t*) computes the temporal derivative of the signal (see Supplementary Note 1), and the tumble rate (Eq. (1)) is λ(*t*) ≃ 1 − *βdS*/*dt*, so that the drift velocity is given by Eq. (5) (see Sect. 2.3 in Ref.^41^). This result is consistent with fast adaptation dynamics approaching gradient-sensing^31,41^.

Note that the system is in the small response regime (Eq. (7)) where KS is applicable (compare Fig. 3d with Fig. 2d). Yet, our AB numerics show that cells with memory can drift faster than predicted by mere gradient sensing (KS) when navigating environments with spatially correlated fluctuations, thus indicating a role for cellular memory in using spatial information beyond local gradients.

### DERIVATION OF DRIFT SPEED IN RUGGED LANDSCAPES

To capture the numerically observed enhancement of AB chemotaxis in rugged landscapes, we extend de Gennes’ analytical derivation of the drift velocity to incorporate the interaction of memory with the landscape fluctuations. To facilitate our analysis, in the rest of this section we work in the OU limit of the landscape, i.e., *m* → 0 in Eq. (9).

Consider a population of cells navigating the rugged landscape *S_η_*(*x*). Under chemotaxis, the average duration of runs up the gradient 〈*t*^+^〉_AB_ is larger than the duration of runs down the gradient 〈*t*^−^〉_AB_. Following de Gennes^36^, it can be shown that the (non-dimensionalised) average velocity of the cells is:

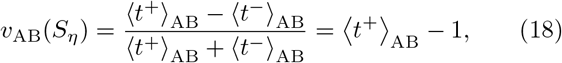

where we use the fact that 〈*t*^+^〉_AB_ + 〈*t*^−^〉_AB_ = 2, i.e., the sum of the average duration of one up-hill and one downhill run is equal to two runs. Equation (18) just states that the average cell velocity is the excess duration of the average up-gradient run beyond the duration of an average run.

Using the ergodicity of the run-and-tumble process, the expectation over AB tra jectories becomes a time integral:

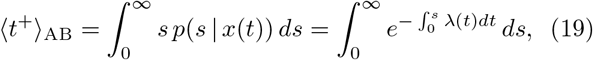

where 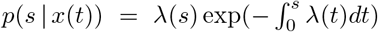 is the conditional probability density of Poisson tumble times given the path *x*(*t*). In the small response regime (Eq. (7)), we can expand the exponential to second order to obtain

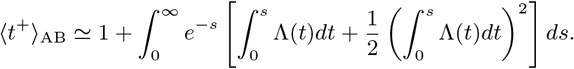

We now depart from de Gennes’ derivation and consider the cell velocity (Eq. (18)) averaged over realisations of the landscape *S_η_*:

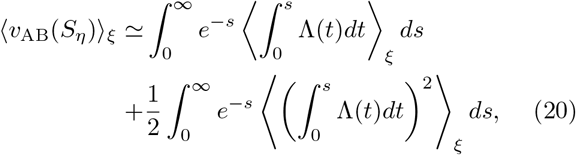

where the first term does not depend on the spatial noise:

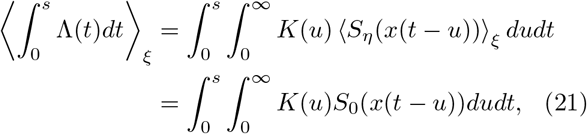

and the second term contains the effect of the spatial correlations as a result of the overlap between the memory kernel and the spatial covariance (Eq. (11)):

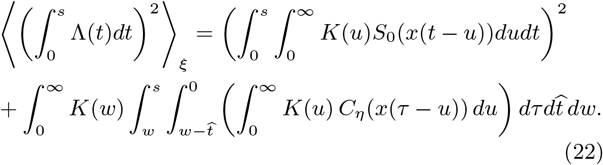

Here 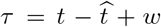 represents the delay between the input *η*^0^(*x*(*t* − *τ*)) and the output Λ(*t*), and the limits of integration reflect causality.

Collating Eqs. (19)–(22) and integrating, we obtain our approximation of the drift velocity in rugged landscapes:

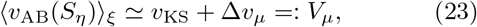

where Eq. (21) gives rise^36^ to the KS drift velocity (Eq. (5)):

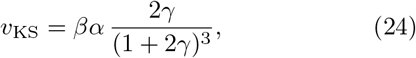

and Eq. (22) leads to the correction due to spatial correlations:

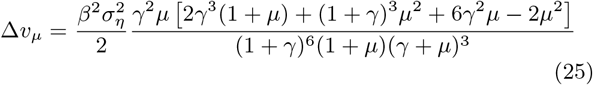

For further details, see Supplementary Note 5.

Our approximation *V_μ_* recovers the KS drift velocity in different limits: for deterministic and uncorrelated landscapes (Δ*v_μ_* → 0 as *μ* → {0, ∞}); in the zero and infinite memory limits (Δ*v_μ_* → 0 as *γ* → {0, ∞}); as well as in the limit of vanishing gradient (*V_μ_* → *v*_KS_ as *α* → 0), since (Eq. (16)) is required to derive Δ*v_μ_*.

### THE EFFECT OF MEMORY ON THE DRIFT SPEED

Our approximation (Eq. (23)) makes explicit the fact that cells use both the local gradient (through *v*_KS_) and the spatial correlations (through Δ*v_μ_*) to navigate rugged landscapes. Fig. 3c shows that, in contrast to the KS drift, our *V_μ_* predicts the enhanced cell velocity in the AB simulations, 〈*v*_AB_(*S_η_*)〉_*ξ*_, and its dependence on the landscape length scale *μ* for a broad range of memory, *γ*.

Fig. 4 compares the predicted maximal velocity and the optimal memory at which it is achieved,

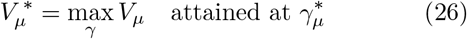

with the numerical simulations. As expected, we recover the KS behaviour in both limits of deterministic (*μ* → ∞) and uncorrelated (*μ* → 0) landscapes, when there is no advantage in using memory to use the statistical correlations of the environment. The optimal memory 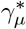 thus emerges as a balance between filtering and tumbling: for 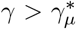, cells improve their noise filtering^32^ but lose orientation due to the larger number of tumbles taken account in their history; on the other hand, for 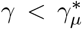, cells are less likely to tumble, but filtering of environmental noise becomes suboptimal. Our results in Fig. 4a show that the velocity of cells with optimal memory is always larger than the gradient-sensing KS drift velocity (i.e., 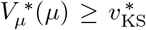, ∀*μ*), with the largest improvement at *μ* ≃ 1/2, the point where the length scale of the environmental correlations are on the order of one half of a run length. As seen in Fig. 4b, the corresponding optimal memory is also always larger than the KS memory: 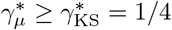. Our calculations show that it is advantageous to increase the memory when the correlations are of the same order as the length of the run (*μ* = 1); yet for correlations longer than one run (*μ* ≥ 1), the presence of random tumblings erode this advantage and the optimal memory returns to the KS value..

**Figure 4.**
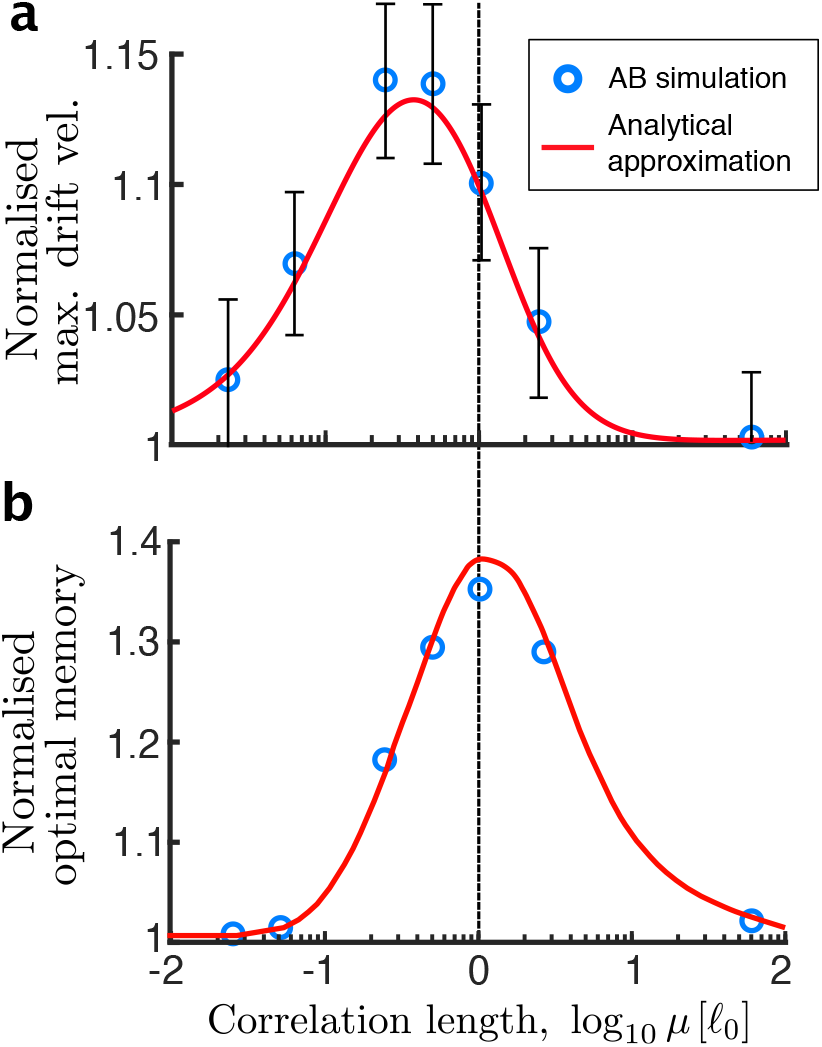
Dependence of the maximal drift velocity and the optimal memory on the environmental correlation length. **a** The maximum drift velocity from the agent-based (AB) numerics at various correlation lengths of the landscape (circles) is well predicted by our approximation (solid line). This maximum, achieved in rugged gradients, exceeds the corresponding deterministic Keller-Segel (KS) drift for all *μ*, and approaches the KS prediction in the white noise (*μ* → 0) and constant gradient (*μ* → ∞) limits. The drift velocities are normalised with respect to the maximum attainable KS drift in deterministic landscapes. The error bars indicate the standard deviation of drift velocity fluctuations across realisations of the rugged landscape at the optimal memory 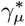 corresponding to this drift speed. **b** The optimal memory against the correlation length shows that the memory at which the drift velocity is maximised is always larger in AB cells than the optimal KS value: 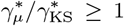. The longest optimal memory for a given correlation length occurs for *μ* = 1 (dotted line), which equals the expected run length. Thus, the tumble rate limits the length scale of perceivable fluctuations.

This behaviour is consistent with models of tethered cells receiving noisy temporal stimuli ^32^. In particular, the mutual information between input and output with a delay *τ* is maximised when maximising the correlation

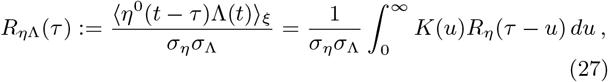

which is a normalised version of the integral in brackets in the second term of Eq. (22). It was shown^32,43^ that Eq. (27) is maximised for a memory corresponding to optimal filtering, and, consistently, our results reflect the importance of noise filtering. For navigation, however, the optimality of filtering is not the only criterion, and it needs to be balanced with the random tumbling time scale which imposes a threshold on the bandwidth of correlations that are useful to improve drift speed.

### THE EFFECT OF MEMORY ON THE HETEROGENEITY OF THE POPULATION

Thus far, we have shown that our approximation predicts well the effect of memory on enhancing the cell velocity in rugged landscapes, although, as seen in Fig. 3c, it overpredicts the velocity of AB cells with short memory 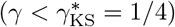 navigating mildly rugged landscapes (0.05 < *μ* < 0.5). The origin of this discrepancy is in the fact that cellular memory has an effect not only on the average cell velocity but also on the heterogeneity of the distribution of AB cells, an effect that is not captured by our approximation in Eq. (23).

To characterise this behaviour further, we carry out additional numerical computations. Intuitively, we expect that cells with short memories will be more sensitive to local irregularities, and hence more prone to becoming disoriented in rugged landscapes. At the population level, this could lead to the appearance of subpopulations of propagating agents. On the other hand, cells with long memory will average their responses over extended patches of the landscape, thus being less sensitive to local fluctuations of the landscape and maintaining the unimodality of the distribution.

A numerical illustration of this behaviour is presented in Fig. 5a, where we show the long-term AB numerics of two population of cells (one with long memories, another with short memories) starting from an initial Gaussian distribution and navigating a rugged landscape *S_η_*. In the KS model, it is known that a Gaussian population remains Gaussian for all times^44^. Indeed, our AB numerics show that when cells have relatively long memories (*γ* = 1), the population does remain unimodal. However, the population of cells with short memories (*γ* = 0.05) goes from being Gaussian to multimodal (i.e., with separate subpopulations), as time elapses. This behaviour is persistent over long times.

**Figure 5.**
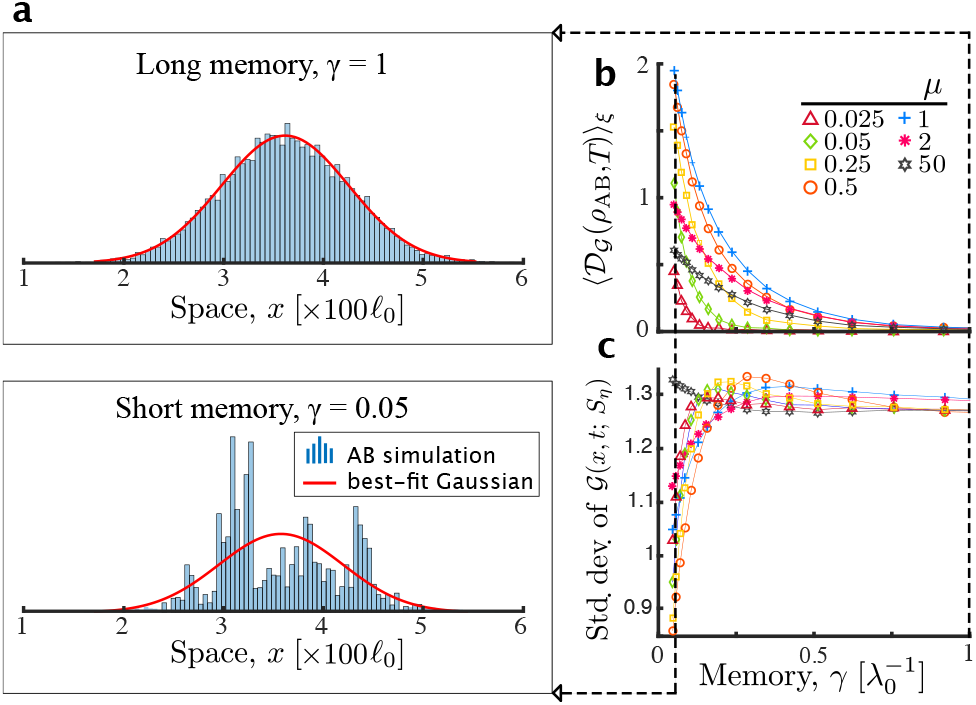
The cellular memory controls the population heterogeneity. **a** Snapshots of the agent-based population density *ρ_AB_*(*x, T; Sη*) in a rugged landscape *S_η_* (*βση* = 10^−3^, *μ* = 1) measured at *T* = 4 × 10^3^ for two values of the memory *γ* (histogram) shown with the best-fit Gaussian curves 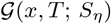 (red line). For short memories (where short depends on the correlation length *μ*) the population loses coherence. **b** As *γ* increases, the population becomes increasingly unimodal Gaussian. This is shown by 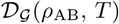, the distance between the population distribution and the best-fit Gaussian (Eq. (28)) averaged over realisations of the landscape, decreasing to zero. **c** For large *γ*, the standard deviation of 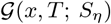 becomes independent of *μ* and *γ*, indicating that randomness arises only from tumbling.

To quantify the loss of unimodality, in Fig. 5b we compute the *L*_2_ distance between the AB distribution *ρ_AB_*(*x, t; S_η_*) and its best Gaussian fit 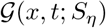 after a long simulation of *T* = 4 × 10^3^:

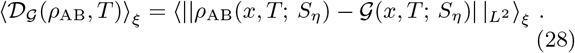

As discussed above, for a rugged landscape with length scale *μ*, the AB distribution becomes increasingly Gaussian for larger *γ*, converging to a Gaussian distribution (Fig. 5b-c). For large *γ*, the standard deviation becomes largely independent of the landscape length scale *μ* (Fig. 5c). This is consistent with the fact that randomness arises not from the landscape but from tumbling, which is common to all cells, thus yielding unimodal population distribution in this limit. When the memory is short, on the other hand, cells use local information and their trajectories depend strongly on the starting positions, leading to distributions far from Gaussian. This dependence on the starting positions is reduced for cells with longer memory, which perceive and average overlapping information of the landscape.

These numerical results suggest that cellular memory could play a role not only in optimising long-term drift velocity (Fig. 4a) but also in controlling population level heterogeneity^45^. A trade-off between both objectives could then allow a more extended exploration of heterogeneous attractant landscapes by the cell population.

## DISCUSSION

Chemotactic navigation relies on the processing of sensory information and its efficient transduction by the cellular mechanisms that control locomotion. Here we studied the run-and-tumble motion of *E. coli* and investigated the role of the cellular memory inherently to the chemotactic response in determining the ability of cells to navigate rugged chemoattractant landscapes. In contrast to previous work, which focused on the deleterious effect of noise on sensing and swimming^11,34^, our model explores how the correlations in the attractant, as encountered through swimming, can be used to enhance chemotactic performance. To this end, we considered an agent-based (AB) model of swimming cells with a history-dependent tumble rate computed by each cell along its trajectory through a signal-transduction model, and compared it to the classical Keller-Segel (KS) model, in which cells instantaneously align their velocity to the chemoattractant gradient independently of their history. Our results confirm that the KS model accurately predicts the behaviour of the AB population when navigating through constant, shallow chemoattractant gradients^31,42^. However, when the chemoattractant landscape has spatially correlated fluctuations superposed to the gradient, AB cells with short-term memory can exhibit greater drift velocity than predicted by mere gradient-alignment.

Building on our numerical observations from the AB model, we extended work by de Gennes^36^ to derive an analytical approximation of the drift velocity that captures the ability of cells to use spatial correlations. The validity of our derivation hinges upon the assumption that the tumbling rate remains close to the adapted state^25^. We show that this assumption, which is less restrictive than the standard shallow gradient assumption^22,42^, is enough to constrain the relationship between the time scales of internal response and perceived stimulus. Specifically, our results hold for rugged environments in the large signal-to-noise regime, even if the typical shallow gradient assumption breaks down. Since our model only allows small tumble biases, the mechanism described here can be fully attributed to the linear filtering of perceived stimuli, and is therefore distinct from the non-linear response considered in previous work^26,27^. Because our analytic derivation of the drift velocity (Eq. (25)) links population dynamics to cell level parameters, it can be used to guide the experimental design to define parameter regimes where deviations from the Keller-Segel limit are expected.

Our analytical model predicts an enhancement of the drift velocity in rugged environments across a range of correlation lengths and cellular memories (Figs. 3–4). This result is consistent with cells with memory performing a non-local optimisation^28^ beyond local gradient alignment. Importantly, when the landscape fluctuations are negligibly small or they occur on long spatial scales, our model recovers the KS model, so that the best strategy is purely local optimisation, as shown by previous studies^38,46^. Our findings are consistent with our fundamental understanding of bacterial chemotaxis: we find that cells relying only on instantaneous information tumble more often, leading to decreased average run length and drift velocity. Hence there is an ecological benefit for the cell to adjust their memory actively to match the length scales in the environment^13^. We also show that our findings are in agreement with optimal information coding by the chemotactic pathway^32,34^ and provide a link between previous results on memory and filtering in the time-domain^10,32^ to chemotactic spatial navigation.

Our model overpredicts the drift velocity of AB cells with short memory navigating mildly rugged landscapes (Fig. 3c). We showed that, in this regime, suboptimal filtering due to short memory results in the segregation of the bacterial population into long-lived multi-modal distributions (Fig. 5). This numerical observation suggests another distinct role for cellular memory as a means to tune how much the ruggedness of the landscape is reflected in the heterogeneity of the population responses.

Micro-scale ruggedness in the landscape can have diverse origins. Ruggedness can appear naturally as a result of random spatial inhomogeneities associated with porous or particulate media, or due to degrading matter and secretions of other organisms. Such processes generate gradient structures on scales smaller or comparable to the mean run-length of bacteria^1,5^. In these environments, bacteria experience the spreading pulse of chemoattractant as a ruggedness in dilute, shallow background gradients^1,47^. There is evidence for this effect in native *E. coli* environments: in soil, the length scales are estimated to vary between ∼ 2 *μ*m – 1 mm^2^; in the digestive tract, the chemical environment is also thought to have fine-grained spatial structure^3^. Furthermore, some aquatic microbes also exhibit run-tumble motion, albeit at higher run speeds of ∼ 100 *μm* s^−1^ and with longer mean run lengths of 68–346 *μ*m^48^. Studies have found that the habitats of such microbes have inhomogeneities with length scales of ∼ 10 – 1000 *μ*m, commensurate with the run lengths of bacteria. In addition, time-varying spatial ruggedness can also appearasaresult of advection (stirring) ofthe medium, which causes it to form a network of thin, elongated filaments^4,5^ on scales of around 200–1000 *μ*m, again commensurate with the run length of aquatic microbes.

Our one-dimensional model allows us to gain analytical insights into the relationship between memory and environmental fluctuations. In particular, we find that the optimal memory is always smaller than the run length. Hence cells are ‘blind’ to longer correlations in one dimension, a fact that is consistent with short-term memory being typical of preadapted cells^20^. In general, however, bacteria can optimise their adaptive responses (and hence their memory) from seconds to minutes through receptor methylation^13^, suggesting the relevance of longer-scale memory-based sensing. This fact may be linked to the stronger directional persistence of bacterial motion in two or three dimensions. With increased directional persistence, we expect that cells can further improve their drift speed by increasing their memory to exploit positive feedback from the environment during up-gradient runs, i.e., by eliciting large tumble rate reductions in comparison to small changes during down-gradient runs^26,27,29^. However, understanding the interplay between memory and directional persistence would be most relevant when studied in conjunction with an appropriate description of 2D or 3D chemoattractant landscapes For instance, patchy landscapes, which are frequently considered in the ecological theory^49^, can give rise to complex navigation strategies^3^ aimed at optimising the use of memory, swimming velocity and directional persistence. Future theoretical work should therefore move from quantifying chemotactic performance in terms of the drift speed in 1D landscapes towards studying swimming in 2D and 3D heterogeneous landscapes using a generalised performance measure, e.g., the amount of encountered attractant^28^.

Finally, although we have concentrated here on navigation in bacterial chemotaxis, memory-based search strategies are also relevant in other biological contexts for higher animals^28^. At a more conceptual level, other search and exploration optimisation processes could benefit from the consideration of agents with memory as a means to take advantage of spatial correlations. For instance, processes of visual search rely on specific neurons in the early visual cortex with a response function commonly approximated by a bi-lobed function (c.f. Fig. 6 in Ref.^50^). From this perspective, saccadic eye movements could be thought of as a navigation over the image with bi-lobed neurons using their memory to integrate the spatial correlations in the visual field as explored during the visual search. Our work could also serve as inspiration for the development of heuristic methods for optimisation problems over rugged landscapes, where particles with memory could be used to complement gradient methods. Future work would be needed to ascertain the significance of memory for optimisation in high dimensions spaces and its effect in enhancing the efficiency of visual search.

## CODE AVAILABILITY

The code to carry out the simulations and analysis can be found at github.com/barahona-research-group/Chemotaxis-In-Rugged-Landscapes under DOI: 10.5281/zenodo.3365951.

## DATA AVAILABILITY

The data generated during the simulations is available with DOI: 10.14469/hpc/6522.

## ACKNOWLEDGEMENTS

We thank Philipp Thomas, José A. Carrillo and Julius B. Kirkegaard for insightful comments. We thank Eduardo Sontag, on one hand, and Nils Becker and Pieter Rein ten Wolde, on the other, for providing us with their agent-based codes, which helped the development and validation of our models. AG acknowledges funding through a PhD studentship under the BBSRC DTP at Imperial College (BB/M011178/1). MB acknowledges funding from the EPSRC project EP/N014529/1 supporting the EPSRC Centre for Mathematics of Precision Healthcare.

## AUTHOR CONTRIBUTIONS

AG and MB designed the study. AG performed the numerical and analytical work interacting closely with MB. AG and MB wrote the manuscript.

## COMPETING INTERESTS

The authors declare no competing interests.

## Notes

https://github.com/barahona-research-group/Chemotaxis-In-Rugged-Landscapes

https://data.hpc.imperial.ac.uk/resolve/?doi=6522

