## Supplementary Material for "Cellular memory enhances bacterial chemotactic navigation in rugged environments"

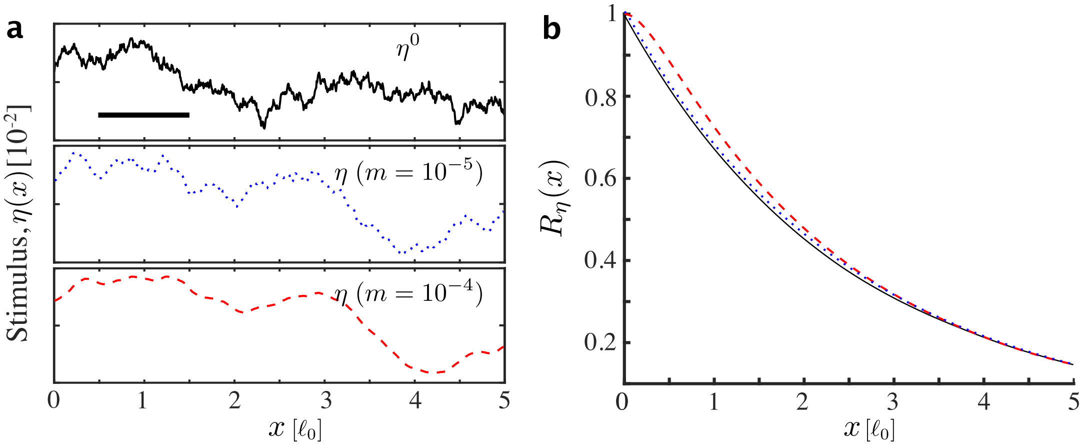


**Supplementary Figure 1: Sample realisations and autocorrelation function of the harmonic oscillator $\eta(x)$ illustrating the deviation from a the limiting Ornstein-Ohlenbeck process $\eta^0(x)$.** **a** Sample trajectories of $\eta^0(x)$ and $\eta(x)$ for correlation length $\mu = 2$ and regularisation factors $m = 10^{-4}$, $10^{-5}$. Bar shows one run length. **b** Autocorrelation function $R_\eta(x) := \langle \eta(x)\eta(x')\rangle_\xi / \sigma_\eta^2$ corresponding to the trajectories in a. The departure from the OU process is not visible for $m < 10^{-5}$. We use the regularised process with $m = 10^{-5}$ for the computation of the agent-based and Keller-Segel models, while performing the analytical computations in the Ornstein-Uhlenbeck limit. The trajectories for $\eta(x)$ were obtained numerically by the Euler-Maruyama scheme with step $\Delta x = 5 \times 10^{-5}$.

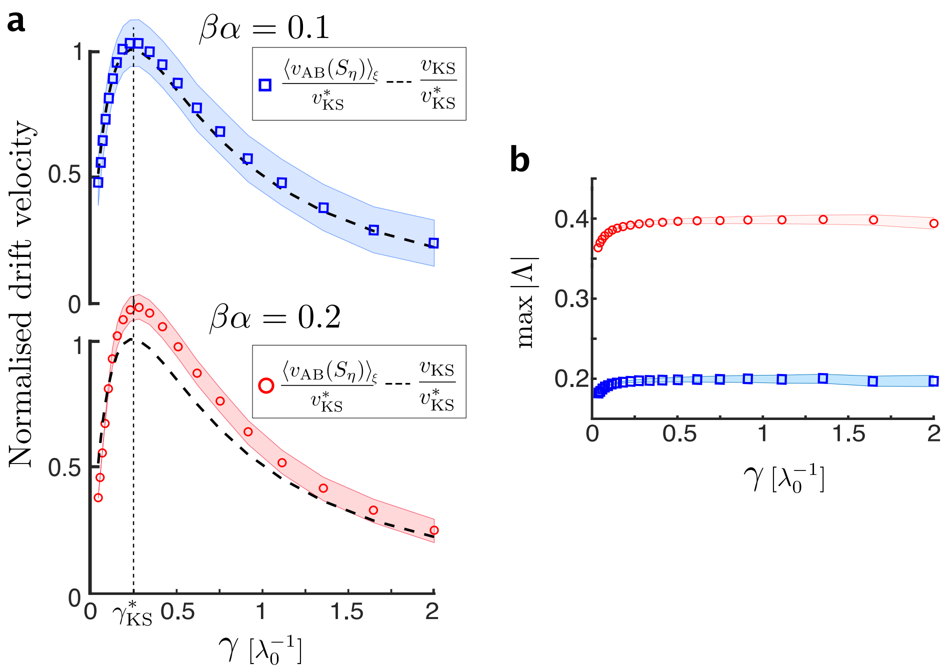


**Supplementary Figure 2: Comparison of agent-based numerics and Keller-Segel approximation in shallow and steep gradients.** **a** For small values of $\beta\alpha$ the Keller-Segel model accurately predicts the drift-velocity of agent-based cells for a large range of memories $\gamma$. For $\beta\alpha > 0.1$ the average velocity of the agents is less well predicted by the Keller-Segel drift velocity. **b** The reason for the discrepancy in a is that the small-response condition $\Lambda \ll 1$ no longer holds for large values of $\beta\alpha$ for all but very small memories $\gamma$.

### Supplementary Note 1

**Tumbling response** Consider cells adapted to the ambient concentration $S(x_0)$ at the beginning of their trajectory $x_0$. The tumbling rate $\lambda(t)$ at time $t$ may then be modelled[1,2] by a linear perturbation depending on the chemoattractant concentrations $S(x(u))$ sensed along the trajectory $x(u)$ in the recent past, $u < t$. Formally,

$$\lambda(t) = 1 - \Lambda(t) \tag{S1}$$

with the *response* $\Lambda(t)$ given by the convolution

$$\Lambda(t) = \int_{-\infty}^{t} K(t-u)S(x(u))\,du. \tag{S2}$$

Here, $K(t)$ is the chemotactic kernel or response function, which for *E. coli* adapting on aspartate has been experimentally characterised[3] as shown on Supplementary Figure 3a. The bi-lobed shape gives positive weighting to recent observations and negative weighting to those further in the past with an exponential decay. As in,[4,5] we consider the dimensionless scaling

$$K(t) = \begin{cases} \frac{\beta}{\gamma}\, e^{-t/\gamma}\left(\frac{t}{\gamma} - \frac{t^2}{2\gamma^2}\right), & t \geq 0 \\ 0, & t < 0 \end{cases} \tag{S3}$$

where, $\gamma$ is the relaxation time (i.e., the cellular *memory*), $\beta$ is the dimensionless signal gain, such that $\beta/\gamma$ dictates the amplitude of the response kernel: $K_{\max} = (\sqrt{2}-1)e^{\sqrt{2}-2}\beta/\gamma$.

The empirical kernel $K(t)$ has several important properties.

1. It is adaptive, that is $\int K(t)dt = 0$.
2. It is causal, i.e. $K(t) = 0$ for $t < 0$. Thus, one may exchange limits in the integral Eq. (S2)
$$\Lambda(t) = \int_{-\infty}^{t} K(t-u)S(x(u))\,du = \int_{0}^{\infty} K(u)S(x(t-u))\,du.$$
3. The kernel computes the instantaneous derivative of the perceived signal $S(t)$ in the limit of vanishing memory $\gamma \to 0$. Indeed, using property 2, we have
$$\begin{aligned}\int_0^\infty K(u)S(t-u)du &= \int_0^\infty K(u)(S(t) - u\frac{dS(t)}{dt} + \mathcal{O}(u^2))du \\ &= \int_0^\infty K(\gamma w)(S(t) - \gamma w\frac{dS(t)}{dt} + \mathcal{O}(\gamma w^2))dw \\ &= \frac{dS(t)}{dt}\int_0^\infty |\gamma K(\gamma w)|dw - \mathcal{O}\left(\gamma \int_0^\infty K(\gamma w)w^2 dw\right) \\ &= \beta\frac{dS(t)}{dt} - \mathcal{O}\left(\gamma \int_0^\infty K(\gamma w)w^2 dw\right),\end{aligned}$$
where in the third equality we used property 1. Taking the limit $\gamma \to 0$ we obtain the result.

**Internal dynamics** The response Eq. (S2) can be obtained from a linear dynamical system with three internal variables $\boldsymbol{y} = (y_0, y_1, y_2)$, as shown by.[5] Let

$$\Lambda(\boldsymbol{y}(t)) = \beta y_2 \tag{S4}$$

be the variable modulating the output. Then, using Eq. (S2),

$$\frac{d}{dt} y_2 = \frac{d}{dt} \int_{-\infty}^{t} \frac{1}{\gamma} e^{-(t-u)/\gamma} \left[ \frac{1}{\gamma}(t-u) - \frac{1}{2\gamma^2}(t-u)^2 \right] S(x(u))\, du\,, \tag{S5}$$

and we may take account of the memory by enlarging the state space to include a finite number of internal degrees of freedom transforming the infinite dimensional integro-differential equation Eq. (S5) into a finite dimensional system of ODEs by the linear chain trick.[6] Repeatedly differentiating Eq. (S5) by Leibniz's rule we obtain

$$\frac{d\boldsymbol{y}}{dt} = \boldsymbol{A}\boldsymbol{y} + \boldsymbol{b}\,, \tag{S6}$$

where

$$\boldsymbol{A} = \begin{pmatrix} -1/\gamma & 0 & 0 \\ 1/\gamma & -1/\gamma & 0 \\ 0 & 2/\gamma & -1/\gamma \end{pmatrix}, \quad \boldsymbol{b} = \begin{pmatrix} \nabla S(x(t)) \cdot v \\ 0 \\ 0 \end{pmatrix}.$$

In this dynamical system, $y_0$ is the input node sensing the changes $S(x(t))$ perceived by a moving cell; $y_2$ is the output node controlling the propensity of tumbling; and $y_1$ is the regulatory node, which acts as a 'buffer' integrating the difference between network response and steady-state output (Supplementary Figure 3a. The unique adapted state (steady-state) of Eq. (S6) is $\boldsymbol{y}_\infty = (0, 0, 0)$, and the eigenvalues of the Jacobian matrix $\boldsymbol{A}$ are negative $(-1/\gamma,\, -1/\gamma,\, -1/\gamma)$, such that $\boldsymbol{y}_\infty$ is locally asymptotically stable. In the following, we will also assume that $\boldsymbol{y}(t)$ remains bounded.

**Agent-based (AB) model** We introduce the agent-based (AB) modelling framework based on Refs.[8,9] At the microscopic level the evolution of the state of a single cell, $t \mapsto (x, v, \boldsymbol{y})$, can be described as

$$\begin{cases} \dfrac{dx}{dt} = v(t)\,, \quad \textit{(position)} \\ \dfrac{d\boldsymbol{y}}{dt} = \boldsymbol{A}\boldsymbol{y} + \boldsymbol{b}\,, \quad \textit{(internal state)} \\ \displaystyle\int_{t_{n-1}}^{t_n} \lambda(\boldsymbol{y}(t))dt = 2\psi_n \quad \textit{(jump times)} \\ v(t) = \nu_n, \quad t \in [t_{n-1}, t_n), \quad \nu_n = -\nu_{n-1} \quad \textit{(velocity-jumps).} \end{cases} \tag{S7}$$

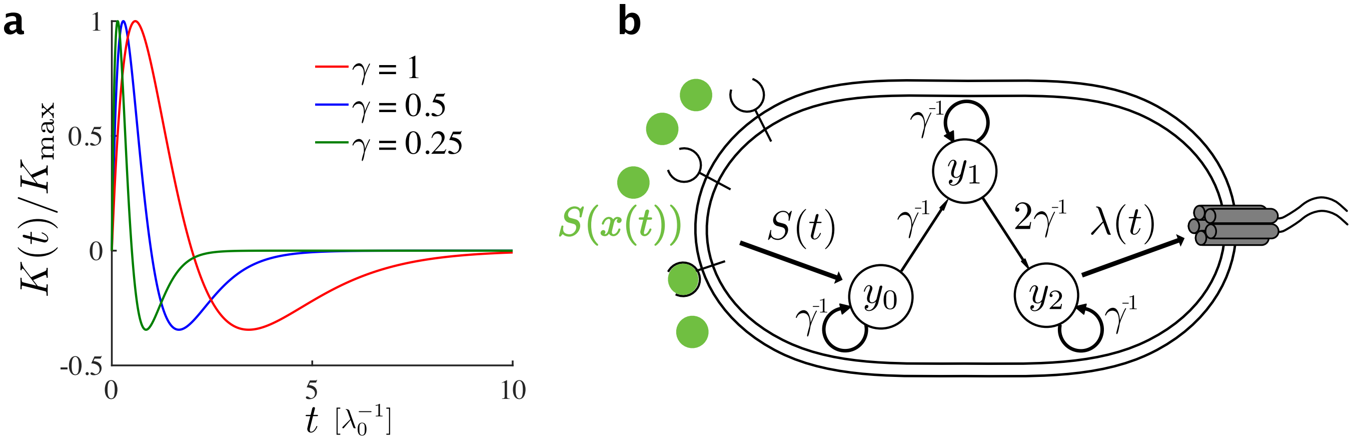


**Supplementary Figure 3: Schematic of the 3-state model of cellular memory and impulse response dynamics.** **a** The normalised response kernel at different values of the memory $\gamma$ and as a function of time **b** Schematic of the 3-state model of cellular memory, whose impulse response is $K(t)$ in a. The extracellular input signal $S(x(t))$ is transduced into a ligand concentration $S(t)$, which modulates the activity of the 'kinase' $y_0$. This kinase relays the signal via an internal 'buffer variable' $y_1$, which in turn produces the response regulator species $y_2$, akin to the CheY protein in the *E. coli* system,[7] in order to control the tumbling rate $\lambda(t)$.

### Supplementary Note 2

**Connection to other modelling frameworks for chemotaxis response** Several other works[1,10] describe the tumbling rate modulations based on the phosphorylation dynamics of the CheY response regulator protein. Although the full model takes account of the non-linear binding kinetics between receptor and chemoattractant molecules, it is typically linearised assuming that the attractant concentration saturates the active receptor population and does not bind to the inactive receptor population. Under such conditions,[10] the tumbling rate is given by:

$$\lambda(t) = \lambda_0 - \beta F(t),$$

with the internal state $F$ evolving according to

$$\frac{dF}{dt} = -\frac{1}{\gamma}(F - F_0) + \omega S'(x), \tag{S8}$$

where $\tau$ is the adaptation timescale (memory); $S'(x)$ is the chemoattractant gradient; and the constant $\omega = \pm\nu N$ is the product of the running velocity $\nu$ and the receptor gain $N$.

It is easy to show that the tumbling rate can then be equivalently expressed in the form of Eqs. (S1)–(S2). Indeed, integrating Eq. (S8) by parts, one obtains

$$
\begin{aligned}
F &= F_0 + \omega e^{-t/\gamma}\left(\left[e^{u/\gamma}S(u)\right]_0^t - \frac{1}{\gamma}\int_0^t e^{u/\gamma}S(u)\,du\right) \\
&= F_0 + \omega e^{-t/\gamma}S(0) + \omega\int_0^t \left(\delta(t-u) - \frac{1}{\gamma}e^{-(t-u)/\gamma}\right)S(u)\,du \\
&= \omega\int_0^t K(t-u)S(u)\,du,
\end{aligned}
$$

assuming that initially $S(0) = 0$ and the adapted state $F_0 = 0$. This expression is in the required form with

$$K(t) = \delta(t) - (1/\gamma)e^{-t/\gamma}, \tag{S9}$$

where $\delta(t)$ is the Dirac-delta function. Note that the kernel Eq. (S9) consists of a singularity at time $t = 0$ combined with a single decaying exponential. In contrast, the kernel we consider in our work starts at the origin and is characterised by a finite excitation time and finite adaptation time leading to one negative overshoot and one positive overshoot.

### Supplementary Note 3

**Derivation of the Keller-Segel drift velocity based on local gradient alignment** In this section, we follow the derivation of the Keller-Segel drift velocity by deGennes.[2] However, instead of a constant gradient, we consider a constant gradient with additive noise, modelled by $S_\eta$. In the rest of this section, we work in the OU limit ($m \to 0$).

Drift velocity is generated when the average runs up the gradients are longer than those the way down. The dimensionless drift velocity on ballistic timescales is therefore

$$v_{\mathrm{AB}}(S_\eta) \simeq \frac{\langle t^+\rangle_{\mathrm{AB}} - \langle t^-\rangle_{\mathrm{AB}}}{\langle t^+\rangle_{\mathrm{AB}} + \langle t^-\rangle_{\mathrm{AB}}} = \langle t^+\rangle_{\mathrm{AB}} - 1, \tag{18}$$

where $t^+$ and $t^-$ are average run times over all possible trajectories for cells moving up and down the gradient for a given realisation of $S_\eta$. As described in the main text, the drift velocity averaged over realisations of the landscape $S_\eta$ becomes

$$\langle v_{\mathrm{AB}}(S_\eta)\rangle_\xi \simeq \int_0^\infty e^{-s}\left\langle \int_0^s \Lambda(t)dt\right\rangle_\xi ds + \frac{1}{2}\int_0^\infty e^{-s}\left\langle\left(\int_0^s \Lambda(t)dt\right)^2\right\rangle_\xi ds. \tag{20}$$

**Remark** Although $|\Lambda(t)| \ll 1$ suffices to justify the expansion Eq. (20), note that it also holds when $\int_0^s \Lambda(t)dt \ll 1$. This is a weaker assumption since $\Lambda(t)$ can be negative causing cancellations to occur in the integral. Also, as argued in the main text, $|\Lambda(t)| \ll 1$ is less restrictive than the classical shallow-gradient assumption of Erban et al.,[11] since it takes account of the internal timescale $\gamma$ in relation to the perceived signal.

In section we describe how to evaluate the first term in Eq. (20), following,[2] while leaving the second term for Supplementary Note 5. Throughout, we will assume that the small response condition holds. In Supplementary Note 4 we derive conditions on $\alpha$, $\beta$, $\sigma_\eta$ that guarantees that it holds, both in constant gradient and in the presence of additive noise.

Then, to obtain the first term, we first compute

$$\left\langle \int_0^s \Lambda(t)dt \right\rangle_\xi = \int_0^s \int_0^\infty K(u)\langle S_\eta(t-u)\rangle_\xi du dt = \int_0^s \int_0^\infty K(u) S_0(x(t-u)) du dt \,. \tag{21}$$

Then, following de Gennes, we use techniques from linear response theory; first, due to the linearity of the response $\Lambda(t)$ we use linear superposition $K(u) = \int_0^\infty K(t)\delta(t-u)du$, and second, assuming that the gradient does not change over a run, we expand $S_0(t-u) = S_0(0) + \frac{dS_0(0)}{dx}(t-u) = S_0(0) + \alpha\,(t-u)$. Then, we have

$$\left\langle \int_0^s \Lambda(t)dt \right\rangle_\xi = \int_0^\infty K(u) \int_0^s \left(S_0(0) + \alpha(t-u)\right) dt du = \alpha \int_0^\infty K(u) \int_0^s (t-u) dt du \,,$$

where in the first equality we used Property 1 of $K(t)$. Further, assuming that the positions before and after the tumble at time $t_n = 0$ are not correlated, one may truncate the kernel $K(t)$ at times $t < u$. Therefore,

$$\left\langle \int_0^s \Lambda(t)dt \right\rangle_\xi \simeq \alpha \int_0^\infty K(u) \int_u^s (t-u) dt du = \frac{\alpha}{2} \int_0^\infty K(u)(s-u)^2 du \,.$$

Inserting into (20) and integrating over $s$ and $u$ yields de Gennes's result[2] for constant gradients

$$\langle v_{\mathrm{AB}}(S_\eta)\rangle_\xi \simeq v_{\mathrm{KS}} = \frac{\alpha}{2} \int_0^\infty K(u) \left( \int_u^\infty e^{-s}(s-u)^2 ds \right) du = \frac{2\beta\alpha\gamma}{(1+2\gamma)^3} \,. \tag{24}$$

Note that no contribution from the spatial randomness appears, since these cancel upon taking averages.

### Supplementary Note 4

**Moment closure** The above derivation of the drift velocity Eq. (S24) relies on the small response assumption

$$|\Lambda(t)| \ll 1, \quad (\textit{small-response}). \tag{7}$$

We now turn to computing the parameter regimes for which this assumption holds, first for constant gradients and then for gradients with additive noise.

**Small-response condition in constant gradients** In constant gradients, the small response condition can be explicitly computed. We compute the output variation over one run:

$$\Lambda(t) = \int_0^\infty K(u)S_0(t-u)\,du = \frac{\beta\alpha}{\gamma}\int_0^\infty e^{-u/\gamma}\left(\frac{u}{\gamma}-\frac{u^2}{2\gamma^2}\right)(t-u)\,du = \beta\alpha\gamma,$$

where we used integration by parts. Hence the amplitude is $|\Lambda(t)| = \beta\alpha\gamma$, and for memories up to $\gamma \sim 1$, we have that the condition

$$\beta\alpha \ll 1 \quad \textit{(shallow perceived gradient)} \tag{S10}$$

is sufficient to ensure that Eq. (7) holds in the main text, i.e., that the cells perceive small variations of the chemoattractant.

**Small-response condition in constant gradients with additive noise** Unlike in the constant gradient case, there is no closed form expression for the response amplitude $|\Lambda|$ when the cell is moving in a random landscape $S_\eta$. Therefore we provide an upper bound as a function of the internal scales $\beta$, $\gamma$ and the external stimulus $\sigma_\eta$, $\mu$. The results below assume that the noise on the landscape is additive, but not otherwise independent of the stochastic model chosen. The following relies on a concentration argument by bounding the mean and the variance of the response amplitude, $\langle|\Lambda|\rangle_\xi$ and $Var_\xi(|\Lambda|)$, respectively, where the averages $\langle\cdot\rangle_\xi$ are taken over the distribution of $\eta^0$.

We first find an upper bound on the average response, $\langle|\Lambda|\rangle_\xi$ in terms of the constants $\alpha$, $\beta$, $\sigma_\eta^2$. We use the scaled kernel $\widetilde{K}(t)$. Then, by the Cauchy-Schwarz inequality we have:

$$\begin{aligned}
\langle|\Lambda(t)|\rangle_\xi^2 \le \langle|\Lambda(t)|^2\rangle_\xi &= \int_0^\infty\int_0^\infty \widetilde{K}(u)\langle S_{\eta^0}(t-u)S_{\eta^0}(t-v)\rangle_\xi \widetilde{K}(v)dudv \\
&= \left(\int_0^\infty \widetilde{K}(u)\alpha(t-u)du\right)^2 + \int_0^\infty\int_0^\infty \widetilde{K}(u)\langle\eta^0(t-u)\eta^0(t-w)\rangle_\xi \widetilde{K}(w)dudw \\
&= \beta^2(\alpha^2+\sigma_\eta^2\sigma_\Lambda^2)\,,
\end{aligned} \tag{S11}$$

where $\sigma_\Lambda^2$ is the normalised response variance.

In the specific case, when the input is a stationary OU process $\eta^0$, the latter can be explicitly computed,

$$\sigma_\Lambda^2(\Gamma) = \frac{\Gamma(\Gamma+3)}{8(\Gamma+1)^3}\,, \tag{S12}$$

where $\Gamma = \gamma/\mu$, defining the ratio between memory and perceived correlations. The stationary points with respect to $\Gamma$ yield a unique maximum $\gamma_\Lambda^* = (\sqrt{7}-2)\mu$. Eq. (S12) is plotted on Supplementary Figure 4, showing excellent agreement between theory and simulations.

Next, we compute an upper bound on the variance $Var(|\Lambda|)$. Note that

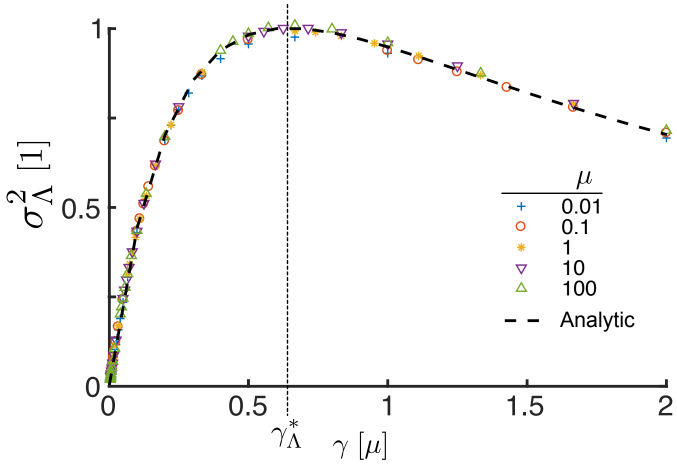


**Supplementary Figure 4: Tumbling rate variance $\sigma^2_\eta$ as a function of $\Gamma = \gamma/\mu$, the memory length $\gamma$ relative to the correlation length $\mu$ of the Ornstein-Uhlenbeck input $\eta^0$.** Symbols show Monte Carlo simulations using the linear dynamical system Eq. (S6) driven by $\eta^0$ at various values of $\mu$. Dashed line shows analytical computations using Eq. (S12). The maximum variance is attained at $\gamma^*_\Lambda = (\sqrt{7}-2)\mu$. The trajectories for $\eta^0$ were obtained numerically using the Euler-Maruyama scheme with step $\Delta x = 10^{-4}$.

$$Var_\xi(|\Lambda|) = \langle|\Lambda|^2\rangle_\xi - \langle|\Lambda|\rangle_\xi^2 = \langle\Lambda^2\rangle_\xi - \langle|\Lambda|\rangle_\xi^2 = Var_\xi(\Lambda) + \langle\Lambda\rangle_\xi^2 - \langle|\Lambda|\rangle_\xi^2 \leq Var_\xi(\Lambda)\,,$$

since $\langle\Lambda\rangle_\xi \leq \langle|\Lambda|\rangle_\xi$. Therefore, it suffices to find a bound on the variance of $\Lambda$. We have

$$\begin{aligned} Var_\xi(\Lambda) &= \langle\Lambda^2\rangle_\xi - \langle\Lambda\rangle_\xi \\ &= \left\langle \int_0^\infty \int_0^\infty K(u)S_\eta(t-u)S_\eta(t-w)\widetilde{K}(w)\,du dw \right\rangle_\xi - \left( \left\langle \int_0^\infty K(u)S_\eta(t-u)du \right\rangle_\xi \right)^2 \\ &= \beta^2\alpha^2 + \int_0^\infty \int_0^\infty K(u)\langle\eta^0(u)\eta^0(w)\rangle_\xi K(w)\,du dw - \beta^2\alpha^2 = \beta^2\sigma^2_\eta\sigma^2_\Lambda\gamma^2\,, \end{aligned} \tag{S13}$$

where we used the stationarity of $\eta^0$ and the definition of $\sigma_\Lambda$ in Eq. (S12).

Collating the above results, we have the following bounds on the mean and variance

$$\langle|\Lambda|\rangle_\xi \leq \beta\sqrt{\alpha^2 + \sigma^2_\eta\sigma^2_\Lambda} \tag{S14}$$

$$Var_\xi(|\Lambda|) \leq \beta^2\sigma^2_\eta\sigma^2_\Lambda\,. \tag{S15}$$

Finally, we use the bounds Eqs. (S14)–(S15) to control the probability of large response amplitudes. By Chebyshev's inequality we obtain

$$\mathbf{P}\left(|\Lambda - \langle\Lambda\rangle_\xi| \geq \epsilon\right) \leq \frac{\beta^2\sigma^2_\eta\sigma^2_\Lambda}{\epsilon^2} \leq \frac{c\beta^2\sigma^2_\eta}{\epsilon^2}\,, \tag{S16}$$

where $c = (10 + 7\sqrt{7})/54$ is the maximum of $\sigma_\Lambda^2$ in Eq. (S12) independently of the system parameters. In the constant gradient limit, as $\mu \to \infty$ or $\sigma_\eta \to 0$, we have $\sigma_\Lambda^2 \to 0$ and hence $P(|\Lambda - \langle\Lambda\rangle_\xi| \geq \epsilon) \to 0$. Thus, $\beta\alpha \ll 1$ is sufficient to control the response, recovering Eq. (S10). Likewise, in the white noise limit, as $\mu \to 0$, we also have $\sigma_\Lambda^2 \to 0$. Therefore, we see that no additional condition on $\sigma_\eta^2$ is necessary for the KS equation to be valid.

However, in general, when the correlation length is not restricted, i.e. $0 < \mu < \infty$, the condition $\beta\alpha \ll 1$ is no longer sufficient for the small response condition to hold, but due to Eq. (S14) and Eq. (S16) additional conditions are needed: $\beta\sqrt{\alpha^2 + c\sigma_\eta^2} = \mathcal{O}(\epsilon)$ and $\sqrt{c}\beta\sigma_\eta \ll \mathcal{O}(\epsilon)$. These are equivalent to requiring

$$\begin{aligned} &\beta\alpha \ll 1 && \textit{(shallow constant gradient)} \\ &\frac{\alpha}{\sigma_\eta} \gg 1 && \textit{(large signal-to-noise ratio)}, \end{aligned} \tag{S17}$$

which means that in addition to Eq. (S10), we require signal-to-noise ratio to be large.

### Supplementary Note 5

**Computation of the second-order contribution** To obtain the contribution from the noise, we begin with

$$\langle v_{\mathrm{AB}}(S_\eta)\rangle_\xi \simeq \int_0^\infty e^{-s}\left\langle \int_0^s \Lambda(t)dt \right\rangle_\xi ds + \frac{1}{2}\int_0^\infty e^{-s}\left\langle \left(\int_0^s \Lambda(t)dt\right)^2 \right\rangle_\xi ds. \tag{20}$$

At this point, we depart from de Gennes' derivation and compute the second-order term to obtain the contribution from the rugged landscape. We proceed by rearranging the integrals

$$\left\langle \left(\int_0^s \Lambda(t)dt\right)^2 \right\rangle_\xi = \left\langle \int_0^s\int_0^s \Lambda(t)\Lambda(\tilde{t})dtd\tilde{t} \right\rangle_\xi = \int_0^s\int_0^s \langle\Lambda(t)\Lambda(\tilde{t})\rangle_\xi\, dtd\tilde{t}\,. \tag{S18}$$

Then, using the definition $S_\eta := S_0 + \eta^0$, we consider the integrand

$$\begin{aligned} \langle\Lambda(t)\Lambda(\tilde{t})\rangle &= \int_0^\infty\int_0^\infty K(w)S_\eta(x(\tilde{t}-w))\langle S_\eta(x(t-u))\rangle_\xi K(u)dudw \\ &= \int_0^\infty\int_0^\infty K(w)S_0(x(\tilde{t}-w))S_0(x(t-u))K(u)\,dudw \\ &\quad + \int_0^\infty\int_0^\infty K(w)\langle\eta^0(x(\tilde{t}-w))\eta^0(x(t-u))\rangle_\xi K(u)\,dudw \\ &= \int_0^\infty\int_0^\infty K(w)S_0(x(t-u))S_0(x(\tilde{t}-w))K(u)\,dudw \\ &\quad + \int_0^\infty K(w)\left(\int_0^\infty K(u)C_\eta(x(\tau-u))\,du\right)dw\,, \end{aligned} \tag{S19}$$

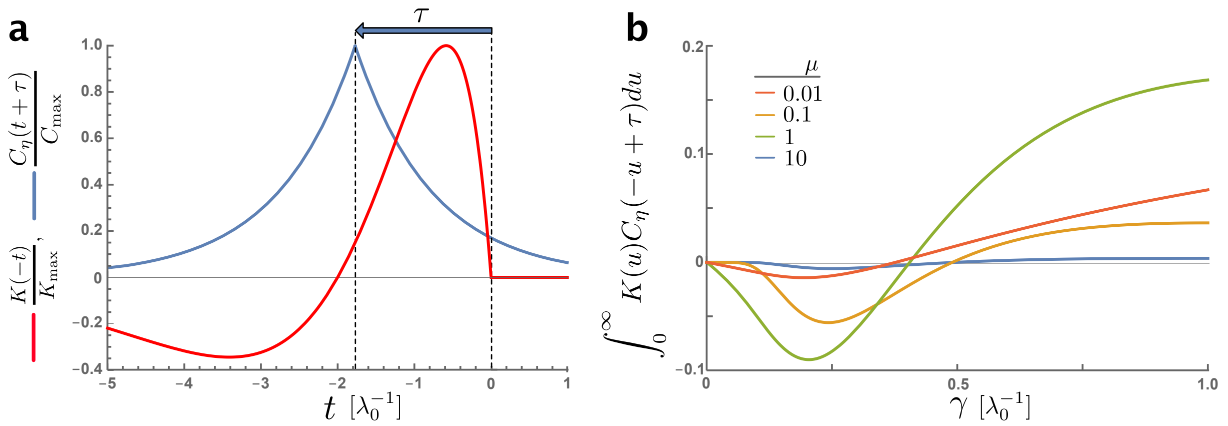


**Supplementary Figure 5: Relationship between the response kernel and the correlation structure of the environment, for finite response time.** **a** Causal responses have a non-negative delay between the input and the output $\tau \geq 0$, which is the case in chemotaxis due to the finite time between sensing the tumbling taking place. This delay results in a time-shift between the response kernel $K(t)$ and the delayed autocorrelation function $C_\eta(t+\tau)$. **b** As a result, for a fixed $\tau$, small memories $\gamma$ lead to sub-optimal filtering. Plotting the overlap integral between the memory kernel and the delayed autocorrelation function as a function $\gamma$ and for different values of the correlation length $\mu$ shows that for a given $\mu$ small $\gamma$ yields a negative contribution to the overlap integral and hence the drift speed (Eq. (S19)).

where the term in brackets defines the overlap integral between the autocovariance function of the input and the chemotactic kernel shifted by $\tau = t - \tilde{t} + w$. This integral is depicted on Supplementary Figure 5 has no contribution in the limits $\mu \to 0, \infty$ since the autocovariance $C_\eta$ is approximately constant and by property 1 of $K(t)$. Between these limits, depending on $\mu$, there is a negative contribution for small memories signifying suboptimal filtering and positive contribution for large memories.

Combining Eqs. (S18)–(S19), we have

$$\left\langle \left( \int_0^s \Lambda(t)dt \right)^2 \right\rangle_\xi = \left( \int_0^s \int_0^\infty K(u) S_0(t-u) du dt \right)^2 + \int_0^\infty K(w) \left( \int_0^s \int_0^s C_{\eta\Lambda}(t-u-\tilde{t}+w)\, d\tilde{t} dt \right) dw \tag{S20}$$

$$= \left( \int_0^s \int_0^\infty K(u) S_0(t-u) du dt \right)^2 + \int_0^\infty K(w) \left( \int_w^s \int_0^{\tilde{t}-w} C_{\eta\Lambda}(t-u-\tilde{t}+w)\, d\tilde{t} dt \right) dw\,. \tag{S21}$$

Note, the first term on the right hand side is the square of Eq. (21), and therefore it is treated as in Supplementary Note 3 to yield $v_{\mathrm{KS}}^2$. In the second term, we first exchanged the integrals, which is allowed

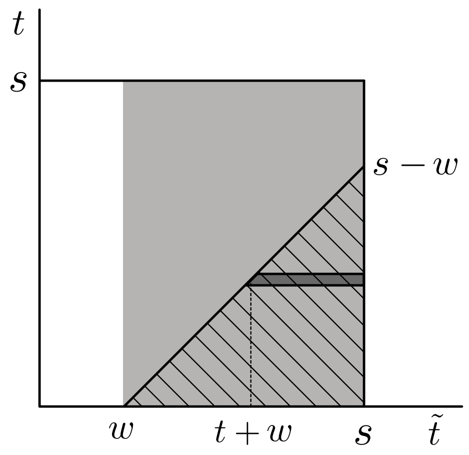


**Supplementary Figure 6: Schematic illustrating the integration region in Eq. (S21) that is consistent with causal responses.** The x and y-axes show the two variables of integration $t$, $\tilde{t}$. Since the swimming direction before and after a tumble are uncorrelated, inputs registered before the tumble (at $t = 0$) do not contribute to the integral. The grey area represents the region of integration to the integral after the tumble. Second, due to causality, the input $\eta^0(\tilde{t} - w)$ must precede the output $\Lambda(t)$. This corresponding integration area is given by the hashed region. Finally, for convenience, we evaluate the double integral by scanning the dark grey area over the hashed region.

since the integrand is bounded due to the exponential decay of $K(t)$, Eq. (S3). Second, as in deGennes' derivation, we ignored the contribution of inputs before the tumbling at time $t = 0$, i.e., $t \in [w, s]$. Thus, information obtained before the tumble does not contribute to the drift speed. Third, due to causality, we need $\tau = t - \tilde{t} + w \geq 0$, i.e., the input at $\eta^0(t - \tau)$ must preceed the output at $\Lambda(t)$. As a result, we only obtain contribution in the region $\tilde{t} \times t \in [w, s] \times [0, \tilde{t} - w]$, as shown by the hashed area in Supplementary Figure 6.

With these assumptions, the correction term due to the random OU fluctuations can be evaluated. We therefore let $\tau = t - \tilde{t} + w$ and consider the input-output covariance in Eq. (S21)

$$\begin{aligned} C_{\eta\Lambda}(\tau) &= \langle \eta^0(t)\Lambda(t+\tau)\rangle_\xi = \int_0^\infty K(t) R_\eta(\tau) dt \\ &= \frac{\beta\sigma_\eta^2}{\gamma}\int_0^\infty \exp\left(-\frac{t}{\gamma} - \frac{|t-\tau|}{\mu}\right)\left(\frac{t}{\gamma} - \frac{t^2}{2\gamma^2}\right) dt \\ &= \beta\sigma_\eta^2 \begin{cases} e^{-\frac{\tau}{\mu}}\frac{\Gamma}{(\Gamma-1)^3} - e^{-\frac{\tau}{\gamma}}\left[\frac{2\Gamma\left(3\Gamma^2+1\right)}{(\Gamma^2-1)^3} - \frac{2\left(\Gamma^2+1\right)}{(\Gamma^2-1)^2}\frac{\tau}{\mu} + \frac{1}{(\Gamma^2-1)}\frac{\tau^2}{\gamma\mu}\right], & \text{if } \tau > 0 \\ \frac{\Gamma}{(\Gamma+1)^3}, & \text{if } \tau = 0 \end{cases} \end{aligned} \tag{S22}$$

where $\Gamma = \gamma/\mu$, as before. Note that although $(\Gamma^2 - 1)$ appears in the denominator, no blow-up occurs when $\Gamma = 1$, as expected since the integrals converge. Combining Eqs. (S21) - (S22) and integrating the second term in Eq. (S21) may be evaluated to yield

$$\left\langle\left(\int_0^s \Lambda(t)dt\right)^2\right\rangle_\xi = \left(\frac{2\beta\alpha\gamma}{(1+2\gamma)^3}\right)^2 + \beta^2\sigma_\eta^2\frac{\gamma^2\mu\left[2\gamma^3(1+\mu)+(1+\gamma)^3\mu^2+6\gamma^2\mu-2\mu^2\right]}{(1+\gamma)^6(1+\mu)(\gamma+\mu)^3} \tag{S23}$$

Finally, collating Eqs. (20), (24) in the main text and (S23), we obtain the drift velocity constant landscapes with additive OU noise keeping only those terms at second order originate from the noise $\eta^0$

$$\langle v_{\mathrm{AB}}(S_\eta)\rangle_\xi := V_\mu \simeq \frac{\alpha\beta\gamma}{(1+2\gamma)^3} + \frac{\beta^2\sigma_\eta^2}{2}\frac{\gamma^2\mu\left[2\gamma^3(1+\mu)+(1+\gamma)^3\mu^2+6\gamma^2\mu-2\mu^2\right]}{(1+\gamma)^6(1+\mu)(\gamma+\mu)^3} = v_{\mathrm{KS}} + \Delta v_\mu\,. \tag{S24}$$
